# Robustly interrogating machine learning-based scoring functions: what are they learning?

**DOI:** 10.1101/2023.10.30.564251

**Authors:** Guy Durant, Fergus Boyles, Kristian Birchall, Brian Marsden, Charlotte M. Deane

## Abstract

**Motivation:** Machine learning-based scoring functions (MLBSFs) have been found to exhibit inconsistent performance on different benchmarks and be prone to learning dataset bias. For the field to develop MLBSFs that learn a generalisable understanding of physics, a more rigorous understanding of how they perform is required.

**Results:** In this work, we compared the performance of a diverse set of popular MLBSFs (RFScore, SIGN, OnionNet-2, Pafnucy, and PointVS) to our proposed baseline models that can only learn dataset biases on a range of benchmarks. We found that these baseline models were competitive in accuracy to these MLBSFs in almost all proposed benchmarks, indicating these models only learn dataset biases. Our tests and provided platform, ToolBoxSF, will enable researchers to robustly interrogate MLBSF performance and determine the effect of dataset biases on their predictions.

**Availability and Implementation:** https://github.com/guydurant/toolboxsf

**Supplementary information:** Supplementary data are available at Bioinformatics online.

## Introduction

Predicting the binding affinity of a protein-ligand complex from its 3D structure has been extensively researched in the past decade (Meli *et al*., 2022). However, doing so accurately and for any protein-ligand complex still poses a significant challenge in computational chemistry (Mobley and Gilson, 2017). Accurately predicting binding affinity would aid in structure-based drug discovery, where the chemical structure of a drug is designed based on the structure of its target, as it would allow design hypotheses to be tested in silico. One proposed methodology, scoring functions (Goodsell *et al*., 1996), which estimate binding affinity based on the features of a single protein-ligand complex structure, offer fast predictions and are suited for high throughput virtual screening and lead optimisation (Bissantz *et al*., 2000). Docking software, such as AutoDock 4 (Morris *et al*., 2009), AutoDock Vina (Trott and Olson, 2010), GOLD (Verdonk *et al*., 2003), and Glide (Friesner *et al*., 2004) commonly use scoring functions to predict the structure of the bound ligand (the pose), its binding affinity and its rank compared to other proposed poses. These scoring functions use either molecular force fields (Huang *et al*., 2006), statistical potentials (Gohlke *et al*., 2000) or linear combinations of empirical terms (Krammer *et al*., 2005). Advancements in machine learning (ML) have enabled the development of ML-based scoring functions (MLBSFs) that appear to outperform other scoring functions in accuracy for predicting binding affinity. Initially, these scoring functions employed classical ML techniques, e.g. tree-based models, and simple features extracted from the protein-ligand complex structure (Ballester and Mitchell, 2010; Wang and Zhang, 2017; Ballester *et al*., 2014; Li *et al*., 2015; Zilian and Sotriffer, 2013; Durrant and McCammon, 2011; Meli *et al*., 2021).

With the emergence of deep learning techniques, scoring functions based on the convolutional neural network (CNN) architecture were built and trained on explicit, voxelised representations of the ligand-protein complex (Francoeur *et al*., 2020) (e.g. Pafnucy (Stepniewska-Dziubinska *et al*., 2018) and KDeep (Jiménez *et al*., 2018)). Newer deep learning methods such as graph neural networks (GNNs) represented atoms as nodes, bonds as edges and used message-passing to pass feature vectors across the graphs to learn higher representations for predicting binding affinity (Moon *et al*., 2022; Karlov *et al*., 2020; Li *et al*., 2021; Scantlebury *et al*., 2023; Volkov *et al*., 2022). Despite the plethora of methods published, there is no clear consensus on which architecture should be used and how to improve scoring function accuracy, given the small differences in performance observed between the methods on the standard benchmarks (Su *et al*., 2019; Carlson et al., 2016).

There have been increasing calls for the field to rethink how we measure MLBSF performance and accuracy (Scantlebury *et al*., 2023; Volkov *et al*., 2022). Most MLBSFs are trained on the PDBBind database (Wang *et al*., 2004), which consists of thousands of protein-ligand complex crystal structures with binding affinity data extracted from the literature. Complexes in CASF 2016, the most popular benchmark for scoring function performance, have very high similarity to data points within the standard training dataset (PDBBind) resulting in an over-optimistic measurement of accuracy as MLBSFs can learn data similarity or “bias” instead of relevant biophysics (Scantlebury *et al*., 2023). Alternative methods of interrogation have been proposed, these include clustered cross-validation (Zhu *et al*., 2022), leave-cluster-out cross-validation (Kramer and Gedeck, 2010), time-splits (Volkov *et al*., 2022) and removing training data similar to the test data (Scantlebury *et al*., 2023; Boyles *et al*., 2020). Unfortunately, due to the widespread use of the CASF 2016 benchmark for evaluating models, researchers can only compare their proposed model to others using that benchmark, exacerbating the problem of inadequate scoring function evaluation. Furthermore, MLBSFs are benchmarked and tested on accurate crystal structures but often will be used for scoring predicted docked ligand poses against non-cognate or predicted structures in a real-world drug discovery setting. These noisy structures are likely to be less accurately predicted compared to the crystal structure, yet this impact has been explored in a limited manner for a few scoring functions (Boyles *et al*., 2022; Scardino *et al*., 2023; Wong et al., 2022; Shen et al., 2021; McNutt *et al*., 2021; Francoeur *et al*., 2020). It has been demonstrated that models trained only on ligand and/or protein identities without explicitly including the interactions between them perform surprisingly well on CASF 2016 (Boyles *et al*., 2020; Volkov et al., 2022). It can be difficult to definitively prove that models are learning bias in the dataset due to the “black box” nature of many machine learning models.

Here we present a platform for interrogating scoring function performance, called ToolBoxSF. First, we reimplemented a diverse set of MLBSFs: RFScore (Ballester and Mitchell, 2010), Pafnucy (Stepniewska-Dziubinska *et al*., 2018), PointVS (Scantlebury *et al*., 2023), SIGN (Li *et al*., 2021) and OnionNet-2 (Wang *et al*., 2021b), to use a consistent API and provide new tests and baseline models to interrogate their performance. We found that simple baseline models trained on only protein and ligand “dataset biases” and unable to learn interactions or the 3D nature of the protein-ligand complex outperformed or had competitive performance to the tested scoring functions in accuracy on a range of benchmarks. We also found that MLBSFs were still partially accurate in nonsensical scenarios like scoring very high root mean squared deviation (RMSD) error poses, ligand docked into a random, wrong protein or complexes with deliberately induced clashes. These results indicate that they are learning dataset bias instead of interpreting the 3D structure. Scoring functions pre-trained or developed for pose classification or ranking did demonstrate sensitivity to clashes though. These results prove that current evaluations of MLBSFs are inadequate at proving usefulness in drug discovery. The provided platform and results should enable researchers to fully and robustly interrogate their models and determine the effect of dataset biases on their predictions.

## Methods

### Training dataset

For consistency, we trained all models on PDBBind 2020 General, the most recent release at the time of writing (Liu *et al*., 2014). It consists of crystal structures of bound protein-ligand complexes with an associated binding affinity label (*K*_*i*_, *K*_*D*_ or *IC*_50_). We excluded complexes that could not be processed by the latest version of RDKit (2023.03.01) (Landrum, 2023) or by OpenBabel (3.1.1) (O’Boyle *et al*., 2011). This left 19079 complexes for training and testing. Structures were prepared as described below (Docking) except the ligand coordinates were not recalculated. In this work, IC_50_, K_*i*_ and K_*D*_ were treated as the same, a common approach in the field (Meli *et al*., 2022) despite the values not being strictly interchangeable (Kalliokoski *et al*., 2013). The pK for each compound was calculated by the following equation:

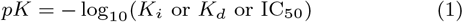

### Docking

Docking was done using Smina, a fork of AutoDock Vina (Koes *et al*., 2013). The default parameters were chosen except “exhaustiveness” (set to 12) and “autobox add” (set to 8Å). The protonation of the ligand and protein were kept consistent with those provided by PDBBind. The MOL2 ligand files provided in PDBBind General 2020 were converted into SDF format for consistency with the docked poses. Their 3D coordinates were recalculated using the ETKDG method from RDKit (Riniker and Landrum, 2015) before docking to ensure the docking software was not able to use the crystal pose to influence its conformational search. Protein files had water molecules and any other non-chain atoms removed by only retaining lines in the PDB file starting with ‘ATOM’.

### Benchmark preparation

To generate a benchmark where ligand bias cannot be used for accurate predictions, the 0 Ligand Bias benchmark, we clustered identical molecules that were bound to different proteins by matching their InChI-Key (Pletnev *et al*., 2012) and took clusters whose mean pK value was within 6 and 7 pK units and whose variance was larger than 1 pK unit. This left 363 complexes as a test set. These two final steps were done to remove identical ligands that had highly similar pK values and to ensure that predicting the mean of the clusters did not artificially increase the accuracy. For example, if two clusters had values concentrated around a low pK and a high pK value respectively, predicting the mean of each cluster would result in a high Pearson’s R between the predicted and true pK values. Peptides, defined as any entry in PDBBind with a ligand code with the letters ‘MER’, were held out to create the Peptides Holdout (2574 complexes). This benchmark tested the scoring functions’ ability to score peptides having never been exposed to them in the training dataset, to be accurate on this benchmark a scoring function must learn an understanding of biophysics that generalises from smaller molecules to peptides. For the 2019 Holdout set, as done in Volkov *et al*. (2022), we took any PDBBind data point with a crystal structure produced from 2019 or later as a test set (1511 complexes). This time split was designed to create a tougher test for scoring functions compared to CASF 2016. To determine the effect of protein structure accuracy on performance, we redocked (Redocked) and crossdocked CASF 2016 ligands into protein conformations that were either bound but with high pocket similarity (CrossDocked (Best)), low pocket similarity (CrossDocked (Worst)), into apo structures (Apo), predicted AlphaFold 2 structures (Alphafold 2) and a random wrong protein (Wrong Protein). The details of how the conformations were picked can be found in the Supplementary Information.

To generate a diverse range of docking errors, we redocked the ligand back into the cognate structure of each protein-ligand complex of CASF 2016, 2019 Holdout and 0 Ligand Bias. We increased the ‘autobox add’ parameter to 20Å and ‘num modes’ to 1000 for Smina. To generate more poses close in accuracy to the true pose, we also minimised the crystal pose using the ‘minimize’ option. Poses were binned by RMSD to the crystal pose using the following ranges: 0-1Å, 1-2Å, 2-4Å, 4-6Å, 6-8Å, 8-10Å, 10-15Å, 15-20Å, 20-25Å and 25-30Å. If available, the pose closest to the mean of the bin was chosen for each test set. To explore the impact of clashes on the models, the crystal pose of the ligand for each complex from CASF 2016, 2019 Holdout and 0 Ligand Bias was progressively translated into the protein 1Å at a time, ten times. To calculate a normal vector for this translation, we normalised the direction vector between the closest ligand and closest protein atom in each protein-ligand complex. Each ligand atom was translated using this normal vector.

### Implementation of scoring functions and models

To compare performance across a range of scoring functions, five popular and diverse models were selected from the literature: RFScore (Ballester and Mitchell, 2010), PointVS (Scantlebury *et al*., 2023), Pafnucy (Stepniewska-Dziubinska *et al*., 2018), SIGN (Li *et al*., 2021) and OnionNet-2 (Wang *et al*., 2021b). RFScore was one of the first methods to use machine learning to predict binding affinity, it uses Random Forest models and counts of protein and ligand elements that are within 12Å of each other. Pafnucy employs a CNN architecture and 3D voxelized representations of the protein-ligand complex. OnionNet-2 also uses a CNN with a 2D image of the counts of each specific amino acid-ligand atom interaction with differing thresholded distances. SIGN and PointVS both use graph neural networks with attention layers for the edges. PointVS is also pre-trained to classify pose accuracy within 2Å and uses this as a prior for its prediction of binding affinity. The pre-trained weights for pose classification of PointVS (Scantlebury *et al*., 2023) were provided by the authors; the model was then fine-tuned using our training set to predict binding affinity. All differences between the original implementations and our modified implementations can be found in the Supplementary Information.

We developed three separate baseline models that represent models that can only learn “bias” in the dataset. All three were developed using tree-based models with architecture and hyperparameters chosen by the FLAML package (Wang *et al*., 2021a) using five-fold cross-validation of the training dataset with CASF 2016 excluded. The LigandBias model is based on the RDKit model from Boyles et al. (Boyles *et al*., 2020), where 1D and 2D descriptors from the RDKit package were calculated to featurise only the ligand. Any descriptor that produced NaN values or extremely large values was excluded, leaving 195 features. The ProteinBias model used counts of each amino acid within the pocket as a feature vector. We defined the protein pocket as any amino acid that had an atom within 15Å of any ligand atom. The impact of this threshold on model performance is explored in the Supplementary Information. This gives the model the identity of the amino acids but not proximity to each other or ligand atoms. Our final model, the BothBias model uses features from both the ProteinBias and LigandBias models. The full information of algorithm and hyperparameters for each baseline model are provided in the Supplementary Information. We also scored all the test sets using Smina (Koes *et al*., 2013) as a baseline for the performance of a non-ML-based scoring function.

The data produced in this work and code for these models have been developed into an easy-to-use platform, called ToolBoxSF, to robustly compare to proposed models from the community and examine if they are learning more than bias. All models have been installed into separate Singularity containers to allow instant and easy use of the models for training or predictions. These wrappers are available on GitHub and as pre-built Singularity containers (https://github.com/guydurant/toolboxsf).

### Metrics

Scoring function accuracy was calculated between their predicted and true values using bootstrapped Pearson R values where data points were sampled with replacement 10000 times to produce 95% confidence intervals.

## Results

### Existing Benchmarks

Evaluation of scoring function accuracy has typically been done using the CASF 2016 benchmark, so we first benchmarked each model on this set (Table 1, CASF 2016). However, the similarity between training (PDBBind) and test set (CASF 2016) makes this an unsuitable benchmark for assessing MLBSF generalisability (Scantlebury *et al*., 2023). We compared five different models that featurise the protein-ligand complex differently: RFScore (Ballester and Mitchell, 2010), Pafnucy (Stepniewska-Dziubinska *et al*., 2018), PointVS (Scantlebury *et al*., 2023), SIGN (Li *et al*., 2021) and OnionNet-2 (Wang *et al*., 2021b). We retrained these scoring functions on our training sets and compared their performance against our baseline models which are unable to learn anything about the structure or interactions of the protein-ligand complex. The model trained on both protein and ligand features that contain no 3D information and so only dataset biases (“BothBias”) had the highest Pearson’s R (0.85) demonstrating that learning biophysics from 3D information is not necessary for close to state-of-the-art performance on the standard CASF 2016 benchmark (0.86) (Wang *et al*., 2021b). High performance on this benchmark has been shown to not be indicative of generalisability (Volkov *et al*., 2022; Scantlebury *et al*., 2023; Zhu *et al*., 2022) but our result goes one step further and demonstrates that even attempting to learn biophysics from structures of the protein-ligand complex provides no additional accuracy. Volkov *et al*. (2022) in an attempt to account for this bias, proposed a time-split where PDBBind data points from 2019 and later were held out as a test set (Table 1 2019 Holdout). BothBias baseline still exceeds the performance of every other model tested except OnionNet-2 with (0.68). This outcome indicates that a time-based split may not be suitable for demonstrating that a scoring function has learnt concepts of biophysics instead of dataset bias. Although both of these benchmarks have value in evaluating the accuracy of scoring functions, it is clear that other benchmarks are required to determine whether a model would be capable of generalising to novel protein or ligand families.

**Table 1.**
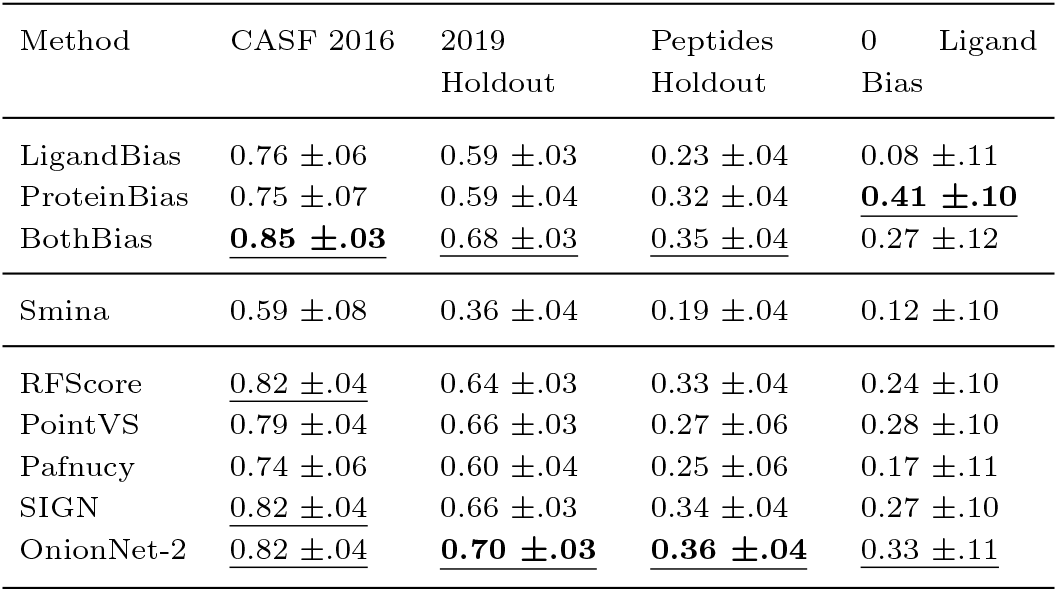
Pearson’s R between predicted and true pK values for protein-ligand complexes for our baseline models (LigandBias, ProteinBias and BothBias), a non-ML-based scoring function (Smina) and five commonly used MLBSFs (RFScore, PointVS, Pafnucy, SIGN and OnionNet-2) on four benchmark datasets (CASF 2016, 2019 Holdout, Peptides Holdout and 0 Ligand Bias). See methods for further details of scoring functions and dataset creation. The highest values are in bold and underlined, second highest is underlined. Error ranges represent the 95% confidence intervals from bootstrapped Pearson’s R (N=10000)

| Method | CASF 2016 | 2019 Holdout | Peptides Holdout | 0 Ligand Bias |
| --- | --- | --- | --- | --- |
| LigandBias | 0.76 ±.06 | 0.59 ±.03 | 0.23 ±.04 | 0.08 ±.11 |
| ProteinBias | 0.75 ±.07 | 0.59 ±.04 | 0.32 ±.04 | **0.41 ±.10** |
| BothBias | **0.85 ±.03** | <u>0.68 ±.03</u> | <u>0.35 ±.04</u> | 0.27 ±.12 |
| Smina | 0.59 ±.08 | 0.36 ±.04 | 0.19 ±.04 | 0.12 ±.10 |
| RFScore | <u>0.82 ±.04</u> | 0.64 ±.03 | 0.33 ±.04 | 0.24 ±.10 |
| PointVS | 0.79 ±.04 | 0.66 ±.03 | 0.27 ±.06 | 0.28 ±.10 |
| Pafnucy | 0.74 ±.06 | 0.60 ±.04 | 0.25 ±.06 | 0.17 ±.11 |
| SIGN | <u>0.82 ±.04</u> | 0.66 ±.03 | 0.34 ±.04 | 0.27 ±.10 |
| OnionNet-2 | <u>0.82 ±.04</u> | **0.70 ±.03** | **0.36 ±.04** | 0.33 ±.11 |

### New Proposed Benchmarks

We propose two benchmarks which evaluate the generalisability of ML scoring functions in different ways. The first utilises the difference between the properties of peptide-protein complexes and ligand-protein complexes found within PDBBind 2020. We removed any peptide-containing complex as a hold-out set from the training dataset. Peptides are difficult to score due to their inherent flexibility and are often much larger than the other ligands in PDBBind (London *et al*., 2010). This makes it a difficult benchmark but success would demonstrate that the models have learnt an understanding of biophysics, such as entropy and changes in solvation, that generalises to peptides. The results in Table 1 show that the BothBias model has similar performance to the best scoring function (OnionNet-2) but neither scores peptides accurately. Our second benchmark takes advantage of scoring functions tending to learn ligand-specific bias in that they are poor at differentiating between the same ligand bound to different proteins (Boyles *et al*., 2020). We identified identical ligands within PDBBind 2020 General that had existed two or more times in the dataset and filtered to ensure these identical ligands’ mean and variance of pKs were centred but spread across the mean pK of the PDBBind dataset (i.e. the training dataset). These groups of identical ligands were then combined into a single set as the 0 Ligand Bias set. On this test set, the BothBias model (0.27) is only as accurate as SIGN (0.27) and is outperformed by OnionNet-2 (0.33) and PointVS (0.28), showing that it is possible to beat this baseline model. However, this is due to learning ligand bias explicitly harming its accuracy, as we can see with the poor performance of the LigandBias model (0.08). The most accurate model is the ProteinBias model (0.41) demonstrating that simply ignoring the ligand entirely out-performs other models currently on this test set. Low performance across all models tested, with Pearson’s R from 0.08 to 0.41, indicates that this is a challenging benchmark. These benchmarks demonstrate that current scoring functions are not able to significantly outperform models trained on bias. Therefore these MLSBFs are both learning bias that does not generalise to this test set and learning little or nothing further.

### Effect of Protein Structure Accuracy on Performance

One deficiency in using the test sets employed above as benchmarks or held-out tests is that they only measure accuracy for scoring crystal structures. Typically scoring functions are used to score docked poses against crystal or predicted structures that might not have an accurate active site conformation for the docked ligand. This introduces noise into the structure as docking predictions may not find the specific interactions or recapture the true binding pose of the crystal structure. To explore the impact of this noise on accuracy, we created alternate docked versions of the CASF 2016 benchmark, which is made up of five structures, each bound to a different ligand, for each of 57 types of proteins (so 285 complexes total) and so contains alternate conformations for the same protein to dock into. We produced six test sets where we re-docked the ligand back into the cognate protein structure (Figure 1 Redocked), cross-docked it into a conformation most similar to its own (Figure 1 Crossdocked (Best)) and again into a conformation most dissimilar (Figure 1 Crossdocked (Worst)). We also docked the ligand into apo (unbound) structures (Figure 1 Apo), predicted AF2 structures (Figure 1 Alphafold 2), and a random protein from CASF 2016 not from its family as a baseline (Figure 1 Wrong Pocket). The differences in structure are shown for a case study (PDB:1E66) in the Supplementary Information to display both the change in 3D structure and how this affects which interactions are formed. These increasingly noisy types of structure demonstrated decreased accuracy when scored by all scoring functions, as shown in Figure 1. The scoring functions were able to retain some accuracy even if the ligand was docked into a completely different protein demonstrating a lower bound of accuracy caused by predictions being dominated by identifying the ligand rather than the nature of the complex. The BothBias model does not appear to be affected as much by the increasing noise as its ligand features are not impacted by changes in conformation and the number of amino acids in the protein pocket does not change significantly across the complex types. These results also suggest as expected that measuring performance on crystal structures provides an upper limit of the ability of scoring functions that is unlikely to be replicated if used in a virtual screen.

**Fig. 1:**
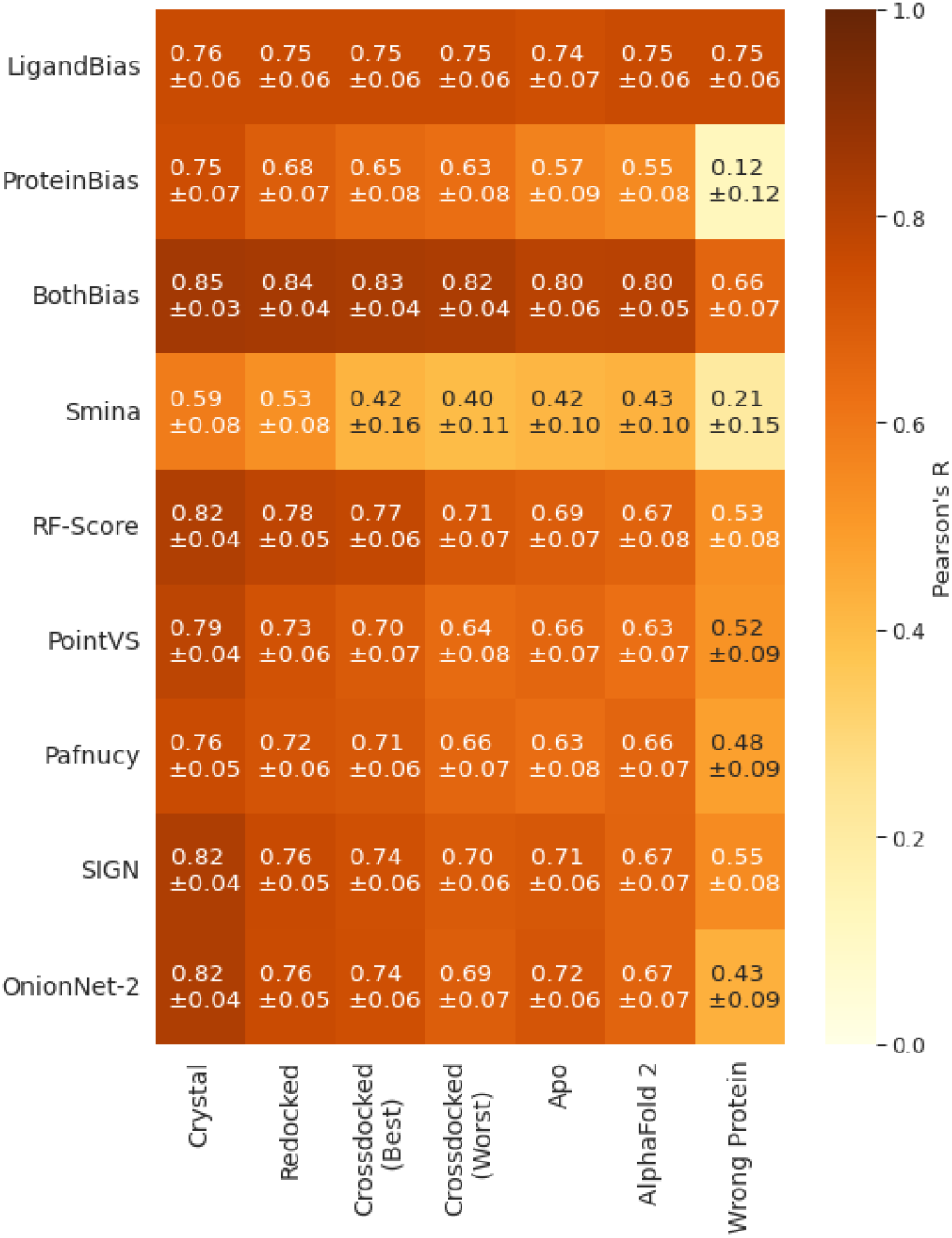
Pearson’s R between predicted and true pK values for protein-ligand complexes for our baseline models (LigandBias, ProteinBias and BothBias), a non-ML-based scoring function (Smina) and five commonly used MLBSFs (RFScore, PointVS, Pafnucy, SIGN and OnionNet-2) on alternate CASF 2016 complex type test sets. Errors are the 95% confidence intervals from the bootstrapped Pearson’s R (N=10000).

### Effect of Docking Accuracy on Performance

To measure the impact of docking accuracy, we considered a diverse set of poses for the CASF 2016 complexes, measured by RMSD. We binned the poses by RMSD first in small ranges (0-1Å and 1-2Å) and then increasingly larger bins with higher inaccuracy to produce 10 test sets. We tested all scoring functions and baseline models but here highlight the results of PointVS, RFScore, SIGN and OnionNet-2 (Figure 2). The results for all other methods can be found in the Supplementary Information. When high-accuracy poses were used, models retained high predictive accuracy relative to scoring the crystal structures when scored by different scoring functions. However, as docking error increased, accuracy decreased and ultimately plateaued at 10Å, except for RFScore which continued to decline beyond this point. Similar to the complex type tests, there was a lower bound for this decrease in performance even at extreme docking errors (25-30Å), where the ligand is no longer bound in the correct site, showing again the models were relying on ligand bias to score protein-ligand complexes. We also explored this effect on 2019 Holdout and 0 Ligand Bias complexes and found the same trend (Supplementary Information). This demonstrates that although docking accuracy is important for binding affinity prediction accuracy, bias is currently a more significant driver of scoring performance as there is still high accuracy for highly inaccurate poses.

**Fig. 2:**
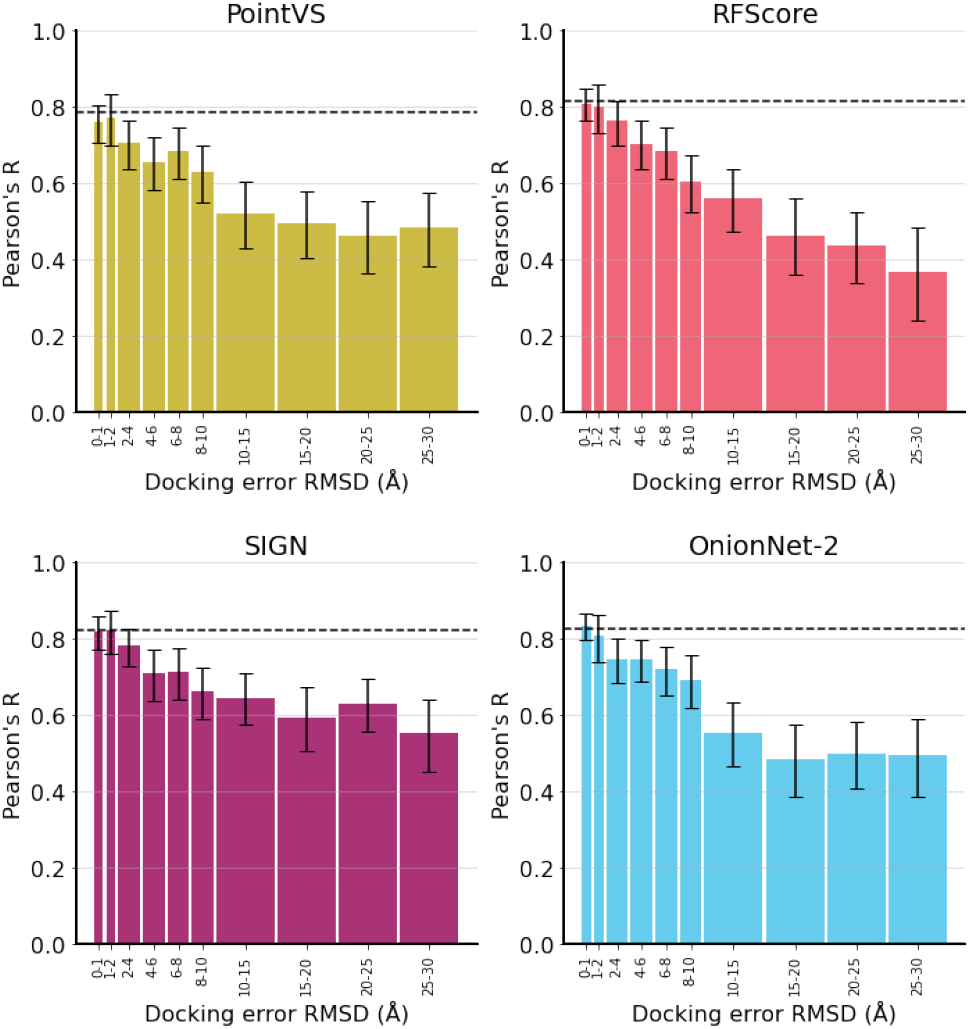
Pearson’s R between predicted and true pK values for protein-ligand complexes for four selected MLBSFs, PointVS, RFScore, SIGN and OnionNet-2, on different accuracy poses of CASF 2016 complexes. Accuracy on the crystal structures of CASF 2016 is shown as a dashed black line. Errors are the 95% confidence intervals from the bootstrapped Pearson’s R (N=10000).

### Clashes

Finally, we investigated scoring function performance when there were clashes in the protein-ligand complex by creating a series of structures where the ligand was translated into the protein for each CASF 2016 complex. Although it is unlikely these scoring functions will come across these types of structures in a drug discovery scenario, the overlap of ligand and protein structure provides such an unrealistic structure with many clashes and few interactions between the protein and ligand. Therefore it is expected that the scoring function should fail to accurately predict binding affinity. The MLBSFs displayed greater sensitivity to translation than the BothBias baseline model; however, most scoring functions displayed only a gradually decreasing performance as the clashes became increasingly severe, again indicating a lower bound (Figure 3). This indicates that the scoring functions only recognise that the ligand is further from the binding site, rather than detecting the unphysical clashes with the protein. The exceptions to these trends are Smina and PointVS, which are both co-trained or pre-trained for pose prediction and demonstrate higher sensitivity to clashes with low or no accuracy on complexes with significant clashes. Again, we also explored this effect on 2019 Holdout and 0 Ligand Bias complexes and found the same trends as for CASF 2016 (Supplementary Information). This suggests that considering pose quality in the training process provides scoring functions with the ability to discriminate between clashing, overlapping structures, and true protein-ligand complex structures.

**Fig. 3:**
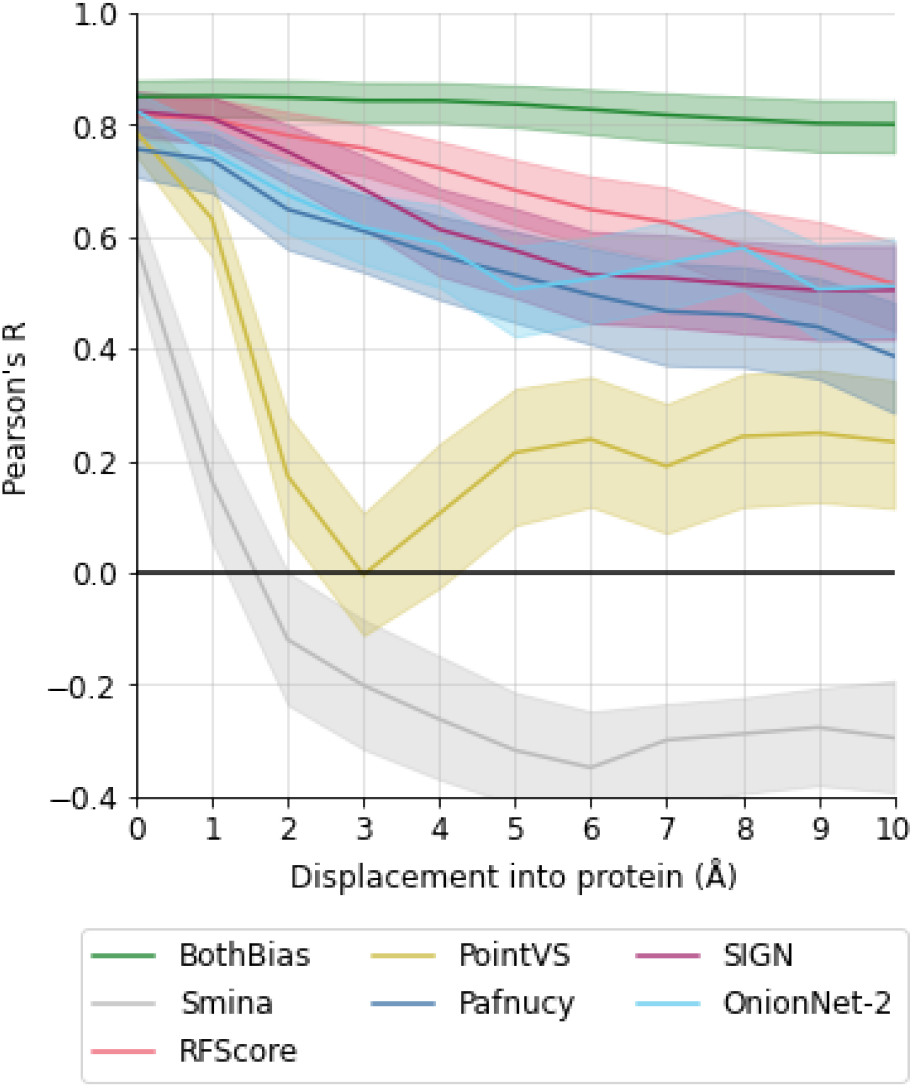
Pearson’s R between predicted and true pK values for protein-ligand complexes for one baseline model (BothBias), a non-ML-based scoring function (Smina) and five commonly used MLBSFs (RFScore, PointVS, Pafnucy, SIGN and OnionNet-2) on progressively displaced ligands into the protein originally from CASF 2016 crystal structures. Errors are the 95% confidence intervals from the bootstrapped Pearson’s R (N=10000).

## Conclusion

In this work, we have demonstrated that state-of-the-art performance on CASF 2016 can be achieved by baseline models using only protein and ligand bias. We propose the 0 Ligand Bias and Peptide Holdout test sets which either explicitly penalise learning ligand bias or require a greater understanding of biophysics, as tougher benchmarks and novel thresholds for improvement from the field. Five popular MLBSFs were equalled or outperformed by baseline models in our tests, indicating that the performance of these scoring functions may be the result of learning dataset bias. We believe our baseline models offer a yardstick for the field as if any proposed scoring function can outperform them, they will have learned more than simple dataset bias.

We examined the effect of noise in the 3D structure of the protein-ligand complex on scoring function performance. The noise introduced by using inaccurate active site conformations or docked poses both resulted in degradation of accuracy in relationship to the amount of noise. This noise being either how dissimilar the active sites are to the cognate crystal structure or the RMSD difference of the pose to the crystal pose. However, both decreases in accuracy had a lower bound showing indifference to the 3D structure input and instead relying on recognising the identity of the ligand.

A further proof that these models are not necessarily learning relevant biophysics is their insensitivity to serious steric clashes between protein and ligands. Translation of the crystal pose into the surface of the protein resulted in a gradual decrease in performance indicating that the scoring functions were only able to recognise that the ligand was further from its true location. The exceptions to this trend, PointVS and Smina, were either pre-trained or developed for pose classification or ranking respectively. These exceptions suggest that scoring functions trained to predict only binding affinity do not learn how sensible a pose is, whilst co-training for another task, such as pose classification, forces it to appreciate clashes.

Overall, this work has provided a meta-analysis of scoring functions and created baseline models that equal existing scoring function accuracy and has provided train-test splits that can help identify if proposed models have learnt more than this simple dataset bias. For the field to progress it will be necessary to design and train models in such a way that they cannot achieve apparent success simply by learning dataset biases. For researchers to prove their proposed scoring functions have learnt more than dataset bias, we have presented rigorous tests and baseline models that can be used for comparisons. All code and dataset splits can be accessed here: (https://github.com/guydurant/toolboxsf).

## Acknowledgments

This work was supported by funding from the Engineering and Physical Sciences Research Council (EPSRC) [grant number EP/S024093/1]. G.D. thanks Dr. Carlos Outeiral for his generation of AlphaFold 2 models for this work.

## Supplementary Information

### 1 Complex Type

#### 1.1 Protein structure selection methodology

For the Redocked set, ligands were simply redocked using Smina back into their cognate structure. As CASF 2016 consists of 5 hand-picked different complex structures with different ligands bound for the same protein type, e.g. HIV protease, for 57 different protein types, there is variability within the conformation for each protein allowing cross-docking into alternative structures within the test set. To generate two classes of cross-docked structures, we aligned the pocket files, provided by PDBBind, using TM-Align (Zhang and Skolnick, 2005) in each set of 5 conformations and calculated TM-Score (Zhang and Skolnick, 2004) for each alignment. The highest TM-Score between a pocket and another pocket within the set was considered the best quality structure for cross-docking (Cross-docked (Best)), and the lowest TM-Score was chosen as the worst quality structure for cross-docking (Cross-docked (Worst)). The original ligand was docked at the site of the cognate ligand for the “best” and “worst” structures.

Apo structures for each cluster were identified by hand from the PDB. The PDB ID for the apo structure and their corresponding CASF 2016 PDB IDs are listed below. Only proteins that had no ligand bound in the active site and had 100% sequence identity for one of the five conformations in the set of the 57 in CASF 2016 were considered. As the conformations in each set in CASF 2016 are not 100% sequence identical, we could not guarantee 100% sequence identical apo structure for each structure in CASF 2016. Of the 57 sets in CASF 2016, only 46 had a suitable apo structure so the final test set was only 230 complexes in size. For each complex, we aligned the apo structure to the original structure and then docked the ligand into the pocket of the apo structure.

For the Alphafold 2 version of CASF 2016, we predicted structures using AlphaFold2 (Jumper *et al*., 2021) for monomers, and AlphafoldMultimer v2.1 (Evans *et al*., 2022) for proteins consisting of multiple polypeptides. Predictions were run as described in the original publication (Jumper *et al*., 2021), including sequences from UniRef (Suzek *et al*., 2007) as well as BFD (Jumper *et al*., 2021) and Mgnify (Mitchell *et al*., 2019). To emulate a realistic blind prediction scenario, we did not include any templates in the prediction, although of course a notable fraction of the targets will have been part of AlphaFold 2’s training set. The ligand was then docked into the aligned AlphaFold structure. All CASF 2016 structures could be successfully predicted, except one: PDB:1YDR. Finally, for the Wrong Protein set, the ligand was docked into a randomly chosen protein not from the same set but still from the CASF 2016 set. We also include visualisations in 3D and 2D how these different complex types impact the docking pose and so the interactions formed for the PDB:1E66.

#### 1.2 PDB IDs of CASF 2016 proteins and their respective Apo PDB ID

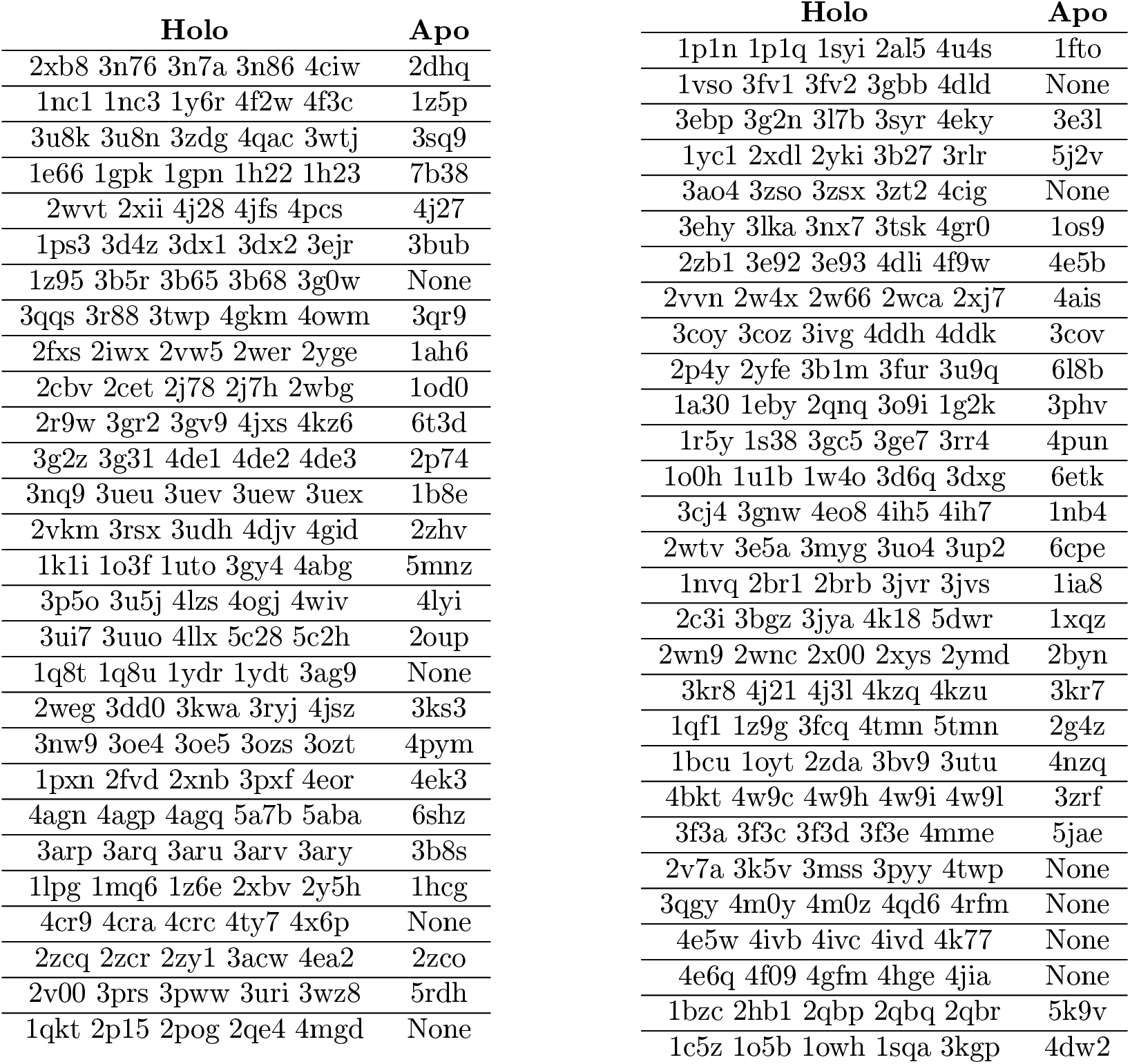

##### 1.3 Visualisation of different complex types for PDB:1E66 and their interactions

**Table 3:**
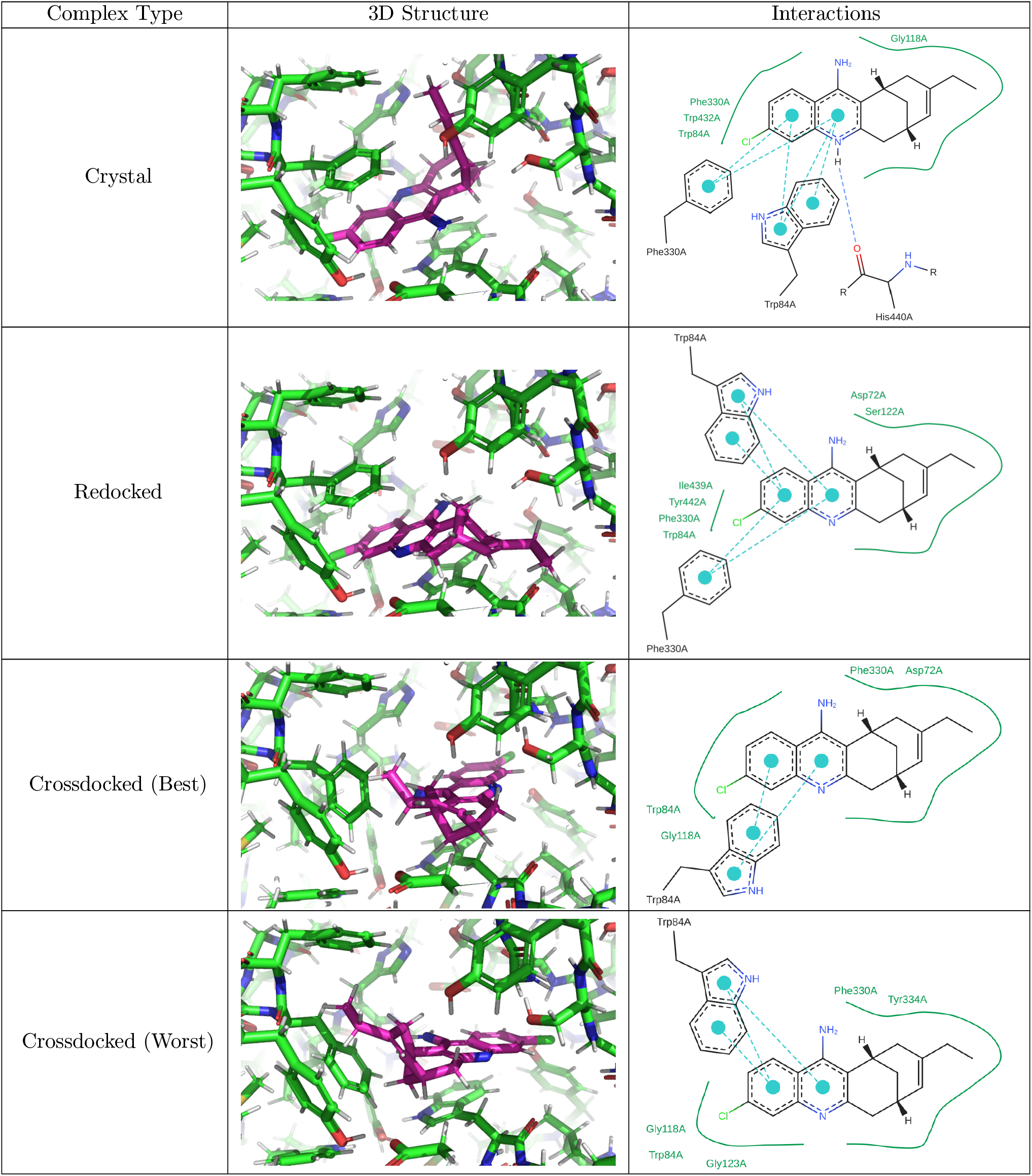
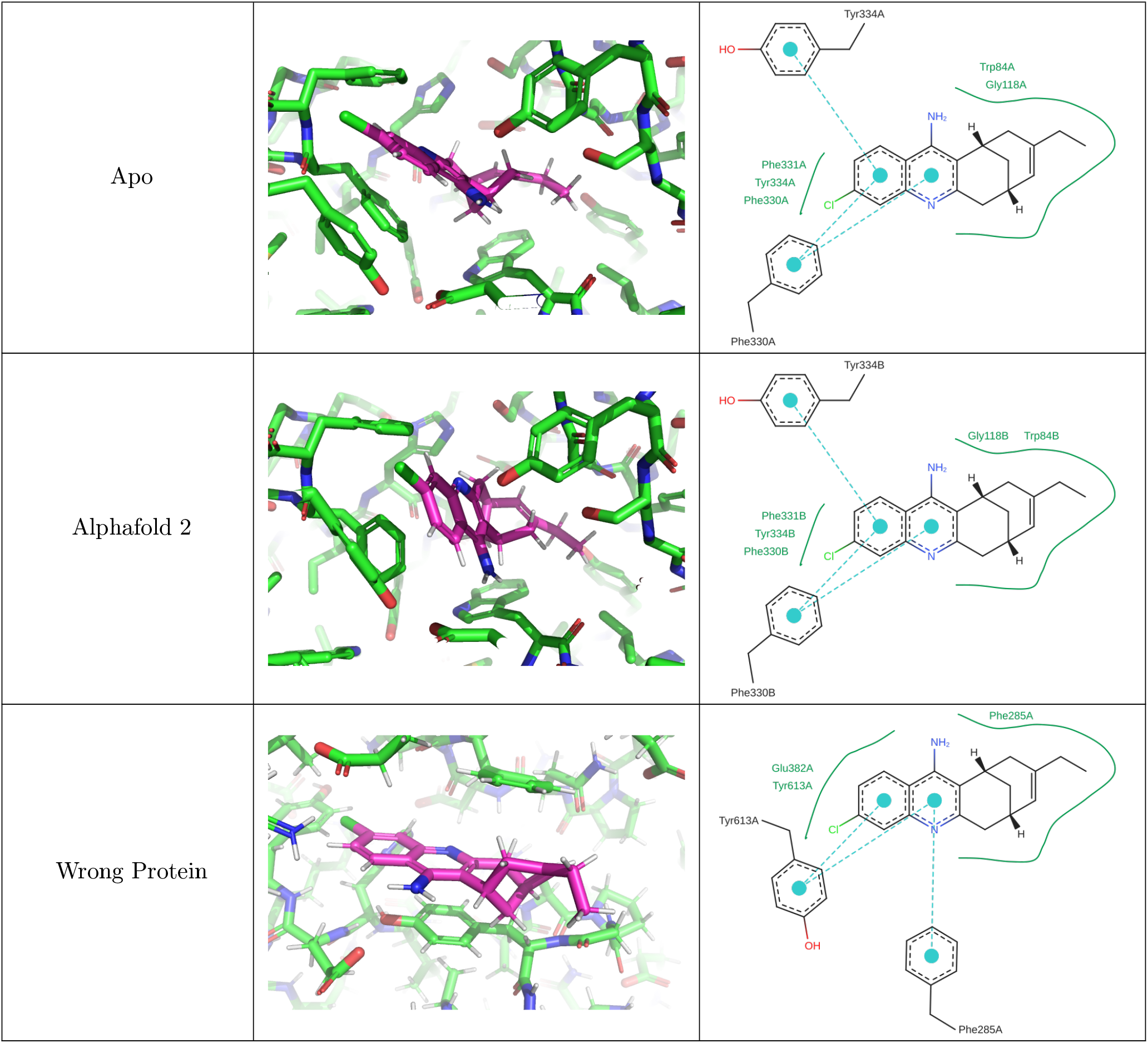
3D and 2D depictions of the protein-ligand complex PDB:1E66 for the different complex types of the Complex Type test sets. The 3D visualisations are at a fixed, consistent orientation and were made using PyMOL Schrödinger, LLC (2015). The 2D depiction of the ligand and the interactions with side chains were generated using PoseEdit Stierand and Rarey (2010) These demonstrate the effect of docking into increasingly noisy protein structures on complex accuracy and the interactions.

### 2 Model implementation

#### 2.1 Machine learning-based scoring function (MLBSF) modified implementations

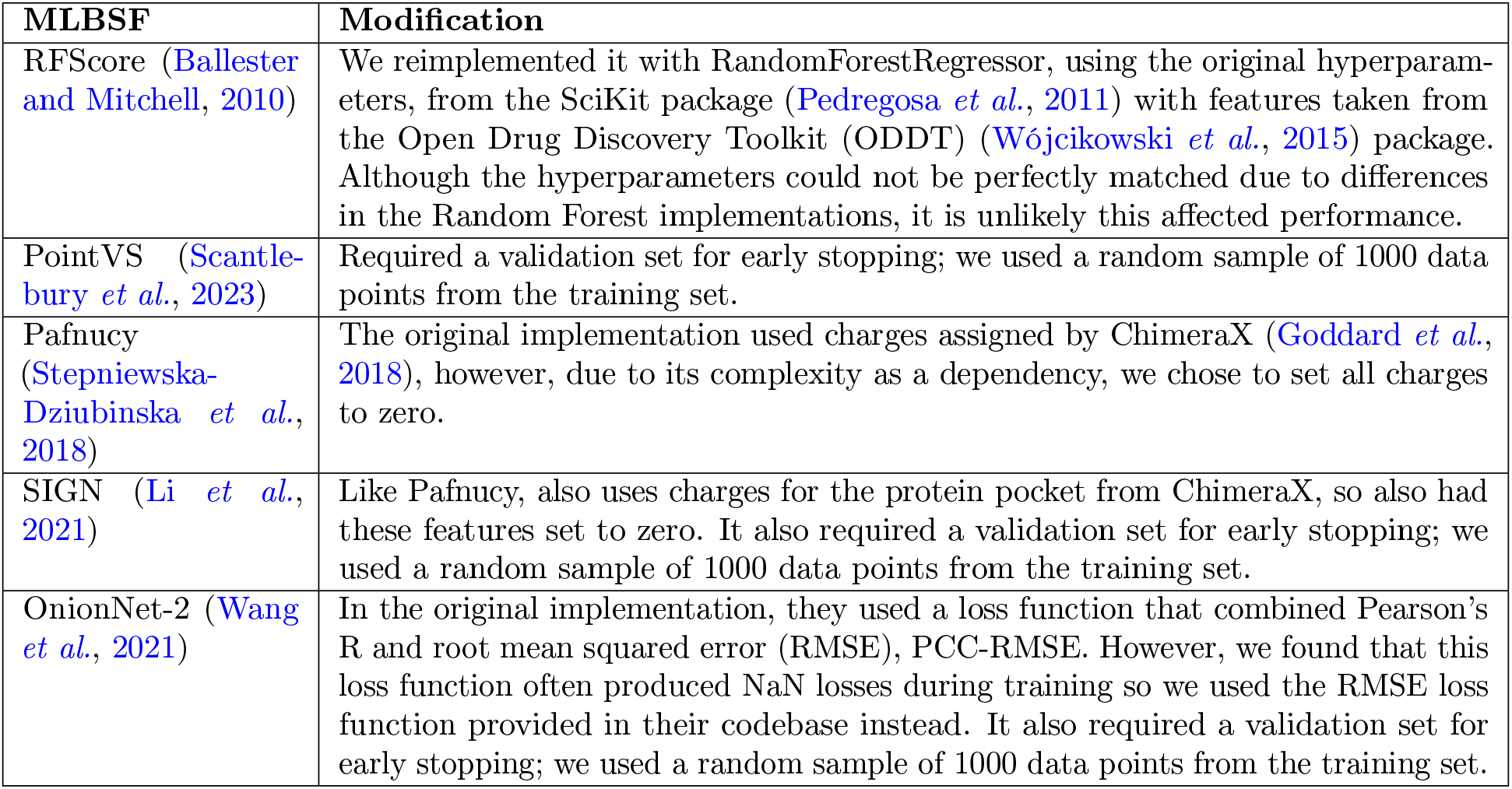

#### 2.2 Hyperparameters of baseline models

| LigandBias | ProteinBias | BothBias |
| --- | --- | --- |
| <u>Random Forest Regressor</u><br>• `n_estimators` = 362<br>• `max_features` = 0.51<br>• `max_leaf_nodes` = 2038 | <u>Random Forest Regressor</u><br>• `n_estimators` = 1447<br>• `max_features` = 0.32<br>• `max_leaf_nodes` = 5460 | <u>LGBMRegressor</u><br>• `n_estimators` = 205<br>• `num_leaves` = 291<br>• `min_child_samples` = 2<br>• `learning_rate` = 0.03<br>• `log_max_bin` = 9<br>• `colsample_bytree` = 0.6<br>• `reg_alpha` = 0.01<br>• `reg_lambda` = 0.01 |

#### 2.3 Impact of protein pocket distance cutoff on accuracy for baseline models

**Figure 1.**
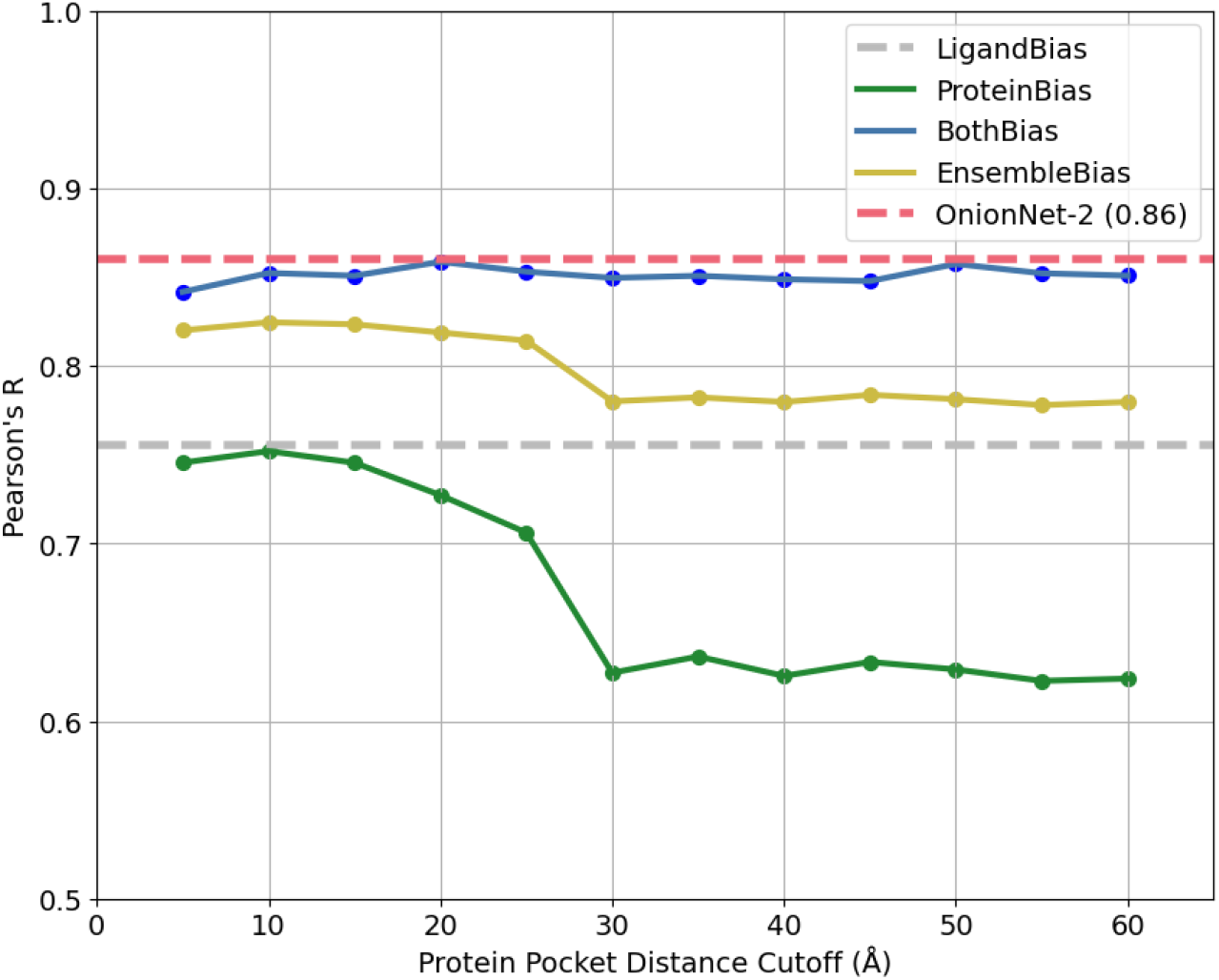
Accuracy of baseline models on the CASF 2016 benchmark when changing the distance threshold, from any protein atom to any ligand atom, to include residues as being part of the protein pocket for featurisation. State-of-the-art performance is included to demonstrate how close the BothBias baseline model is getting to matching it (Pearson’s R = 0.86, OnionNet-2). The EnsembleBias model is the mean prediction of the LigandBias and ProteinBias models. The LigandBias model does not change with distance as it does not use any features derived from the protein of the complex.

### 3 Accuracy of scoring functions and baseline models on protein family hold-out clusters

All the benchmarks investigated in the paper do not reflect a realistic drug discovery scenario as they measure accuracy across many different protein families at once instead of the typical process of screening against a single protein target. We also created the Protein Family Out Benchmarks test sets to measure scoring function accuracy on specific protein families. We clustered the PDBBind dataset, using 90% sequence identity clusters from the PDB (RCBS, 2023) and took any cluster that had more than 100 data points as separate test sets to simulate the screening of a single protein target. The 100 data point size limit ensured there were sufficient data points to evaluate scoring function accuracy. We then trained a new version of each scoring function without each cluster and tested them on the held-out cluster to simulate the scenario of these scoring functions being used on a novel protein target in a virtual screen.

**Figure 2.**
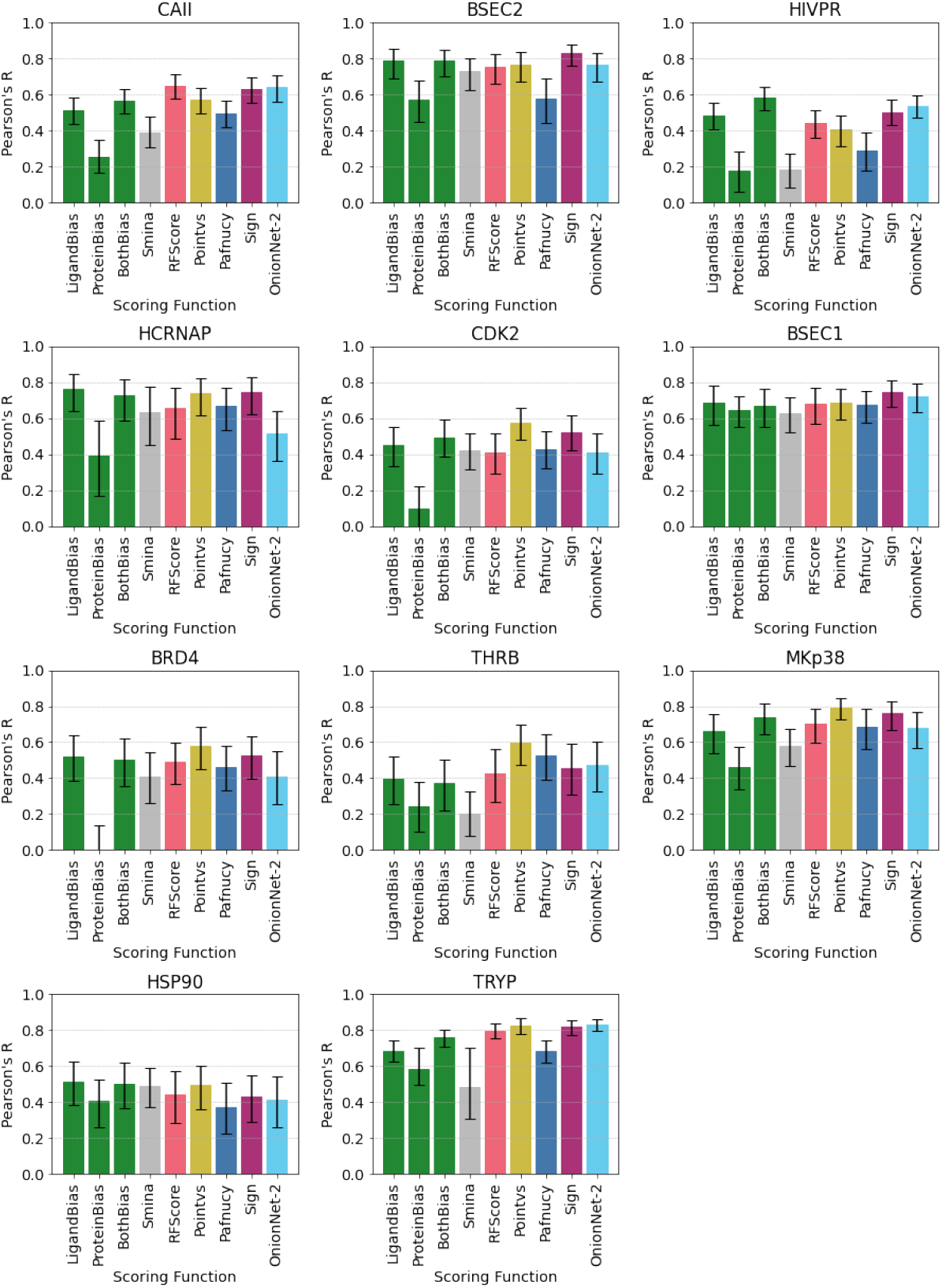
Pearson’s R between predicted and true pK values for protein-ligand complexes for our baseline models (Ligand Bias, Protein Bias and Both Bias), a non-ML-based scoring function (Smina) and five commonly used MLBSFs (RFScore, PointVS, Pafnucy, SIGN and OnionNet-2) for eleven protein family hold-out clusters. These eleven families are Carbonic Anhydrase II (CAII), Beta-secretase (BSEC2), HIV protease (HIVPR), Hepatitis C Virus RNA-polymerase (HCRNAP), Cyclin-dependent kinase 2 (CKD2), Beta-secretase (BSEC1), Bromodomain-containing protein 4 (BRD4), Thrombin (THRB), MAP Kinase p28 (MKp38), Heat Shock Protein 90 (HSP90) and Trypsin (TRYP). Error bars represent the 95% confidence intervals from bootstrapped Pearson’s R (N=10000).

These hold-out tests demonstrate the overall trend that the scoring functions do not vary greatly in the accuracy they achieve, with larger differences in the average ability between protein families than between model accuracy on the same family. The baseline models also follow this trend suggesting that the reason for these differences is probably due to the protein families having different similarities to the training dataset rather than any deeper insight into the scoring functions’ performance. However, some scoring functions can outperform the baseline models in certain families such as CAII, where RFScore and OnionNet-2 (0.64) outperform the BothBias model (0.57). Due to the small differences between the baseline models and the scoring functions and their consistent performance across methods, it is likely that a large amount of accuracy for all methods is due to learning the same bias in the dataset.

### 4 Further scoring functions and baseline models on differing docking accuracy versions of CASF 2016

**Figure 3.**
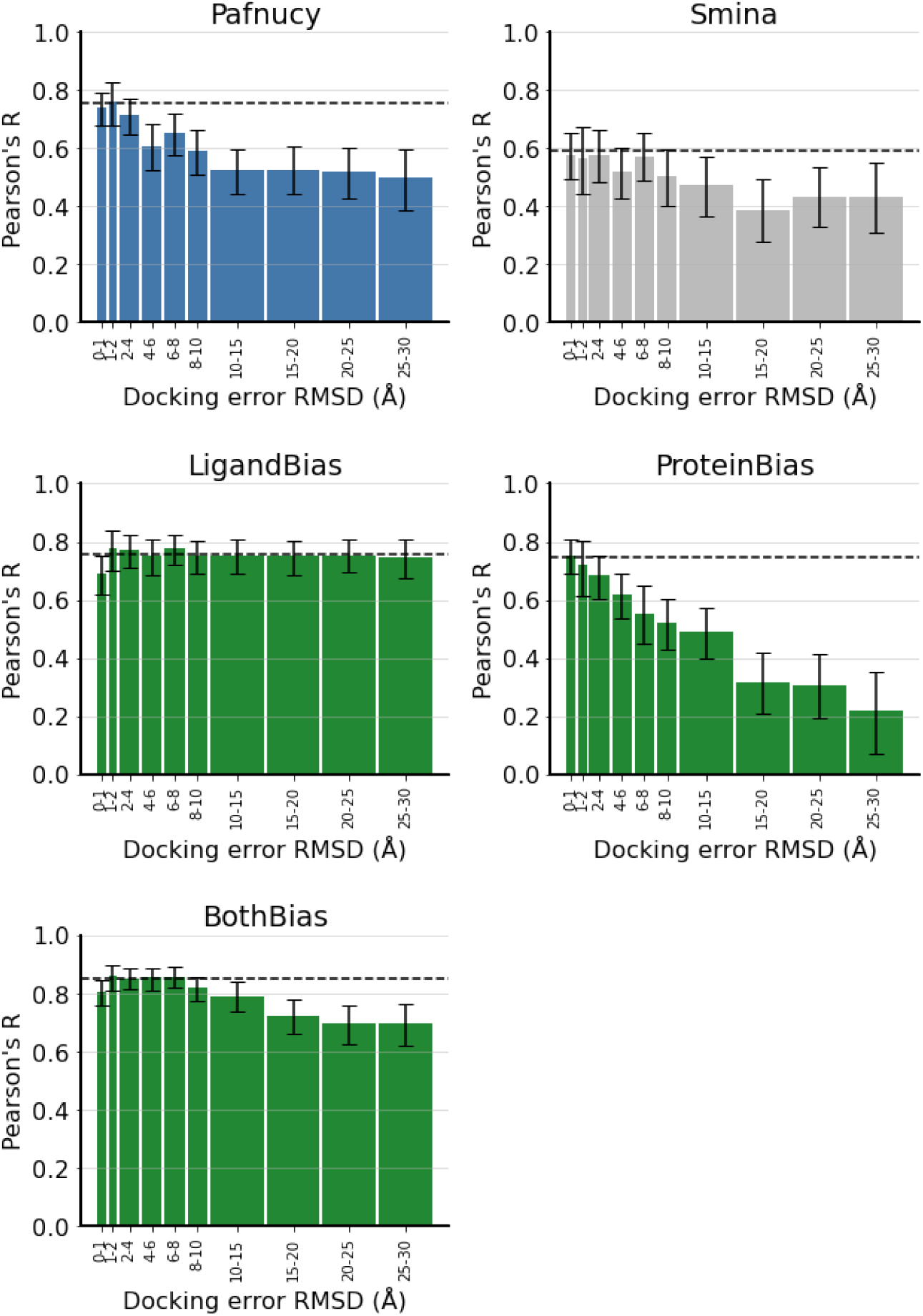
Pearson’s R between predicted and true pK values for protein-ligand complexes for our baseline models (LigandBias, ProteinBias and BothBias), Pafnucy and Smina on different accuracy poses of CASF 2016 complexes. Accuracy on the crystal structures of CASF 2016 is shown as a dashed black line. Errors are the 95% confidence intervals from the bootstrapped Pearson’s R (N=10000).

### 5 Accuracy of scoring functions and baseline models on differing docking accuracy versions of 2019 Holdout

**Figure 4.**
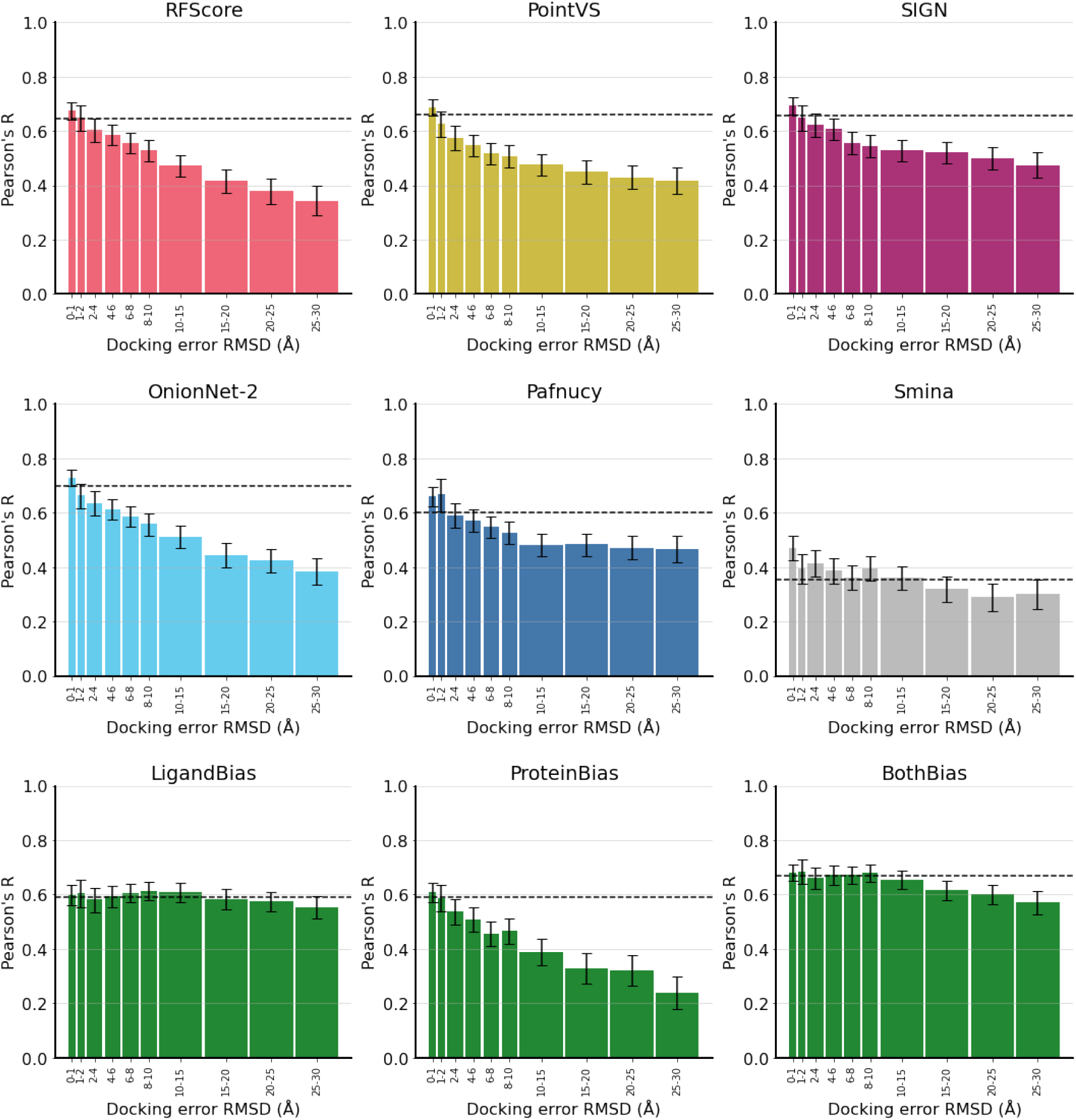
Pearson’s R between predicted and true pK values for protein-ligand complexes for our baseline models (LigandBias, ProteinBias and BothBias), a non-ML-based scoring function (Smina) and five commonly used MLBSFs (RFScore, PointVS, Pafnucy, SIGN and OnionNet-2), on different accuracy poses of 2019 Holdout complexes. Accuracy on the crystal structures of 2019 Holdout is shown as a dashed black line. Errors are the 95% confidence intervals from the bootstrapped Pearson’s R (N=10000).

### 6 Accuracy of scoring functions and baseline models on differing docking accuracy versions of 0 Ligand Bias

**Figure 5.**
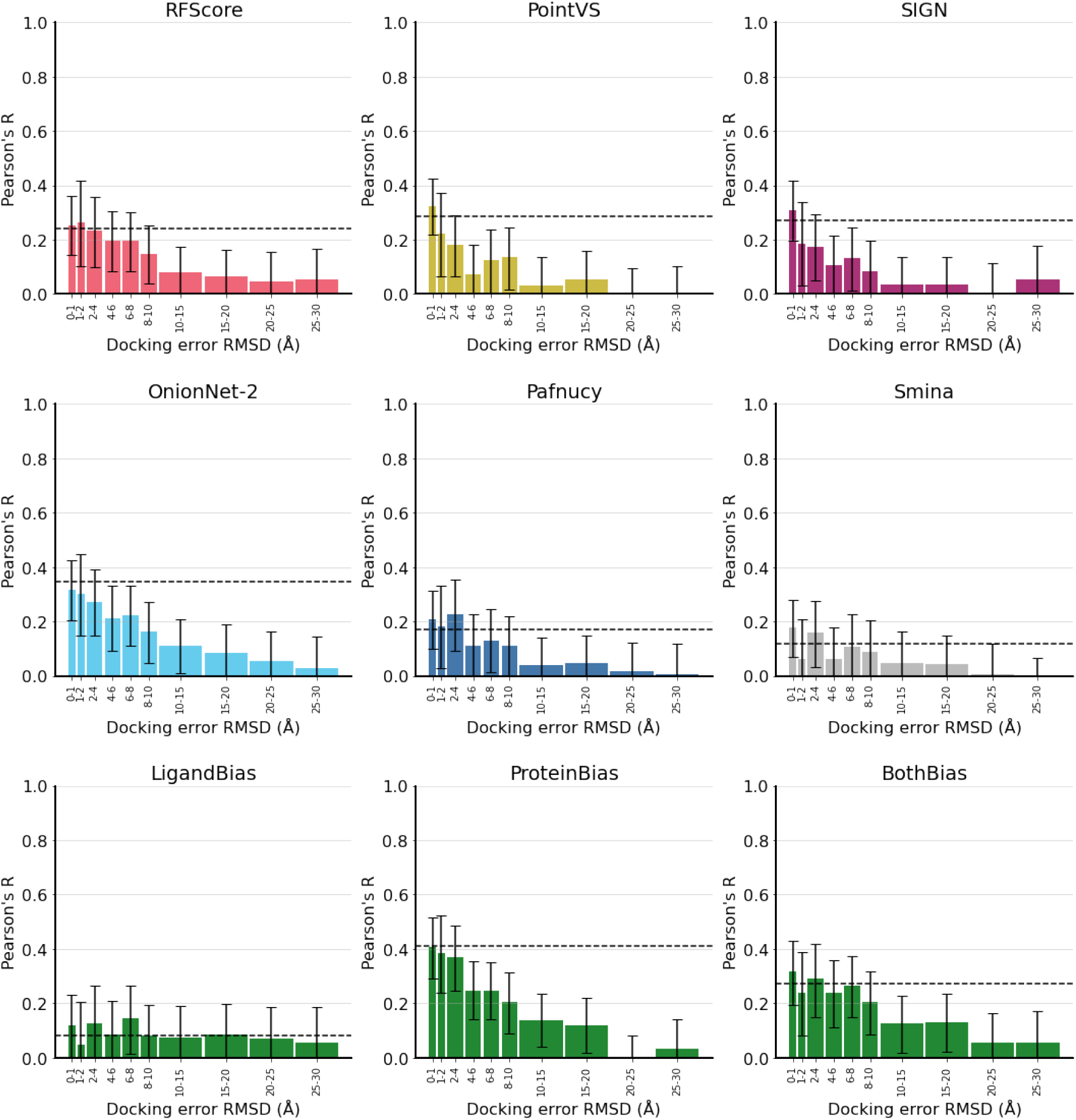
Pearson’s R between predicted and true pK values for protein-ligand complexes for our baseline models (LigandBias, ProteinBias and BothBias), a non-ML-based scoring function (Smina) and five commonly used MLBSFs (RFScore, PointVS, Pafnucy, SIGN and OnionNet-2), on different accuracy poses of 0 Ligand Bias complexes. Accuracy on the crystal structures of 0 Ligand Bias is shown as a dashed black line. Errors are the 95% confidence intervals from the bootstrapped Pearson’s R (N=10000).

### 7 Accuracy of scoring functions and baseline models on progressively displaced ligands of 2019 Holdout

**Figure 6.**
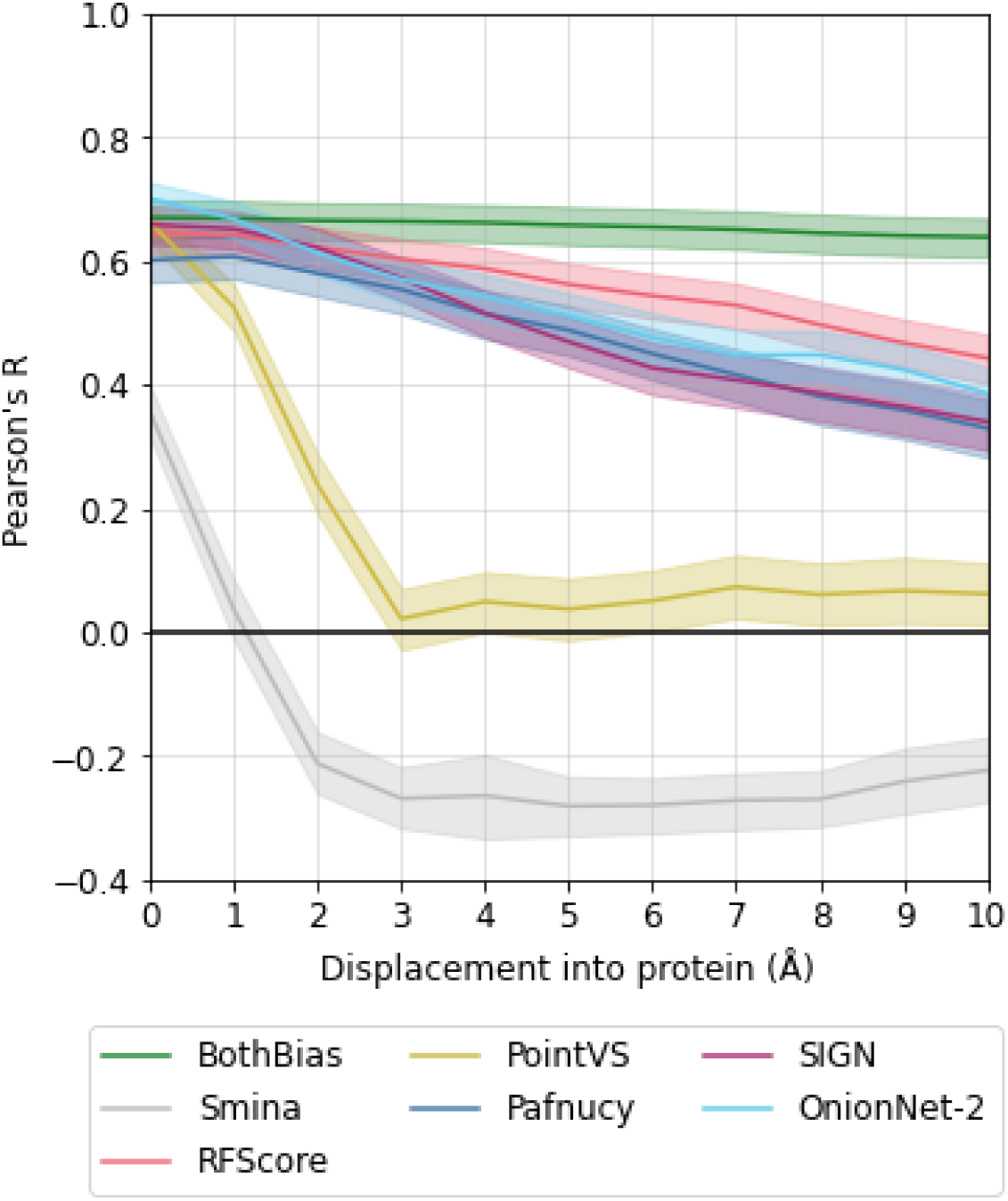
Pearson’s R between predicted and true pK values for protein-ligand complexes for our baseline models (LigandBias, ProteinBias and BothBias), a non-ML-based scoring function (Smina) and five commonly used MLBSFs (RFScore, PointVS, Pafnucy, SIGN and OnionNet-2), on progressively displaced ligands into the protein originally from 2019 Holdout crystal structures. Errors are the 95% confidence intervals from the bootstrapped Pearson’s R (N=10000).

### 8 Accuracy of scoring functions and baseline models on progressively displaced ligands of 0 Ligand Bias

**Figure 7.**
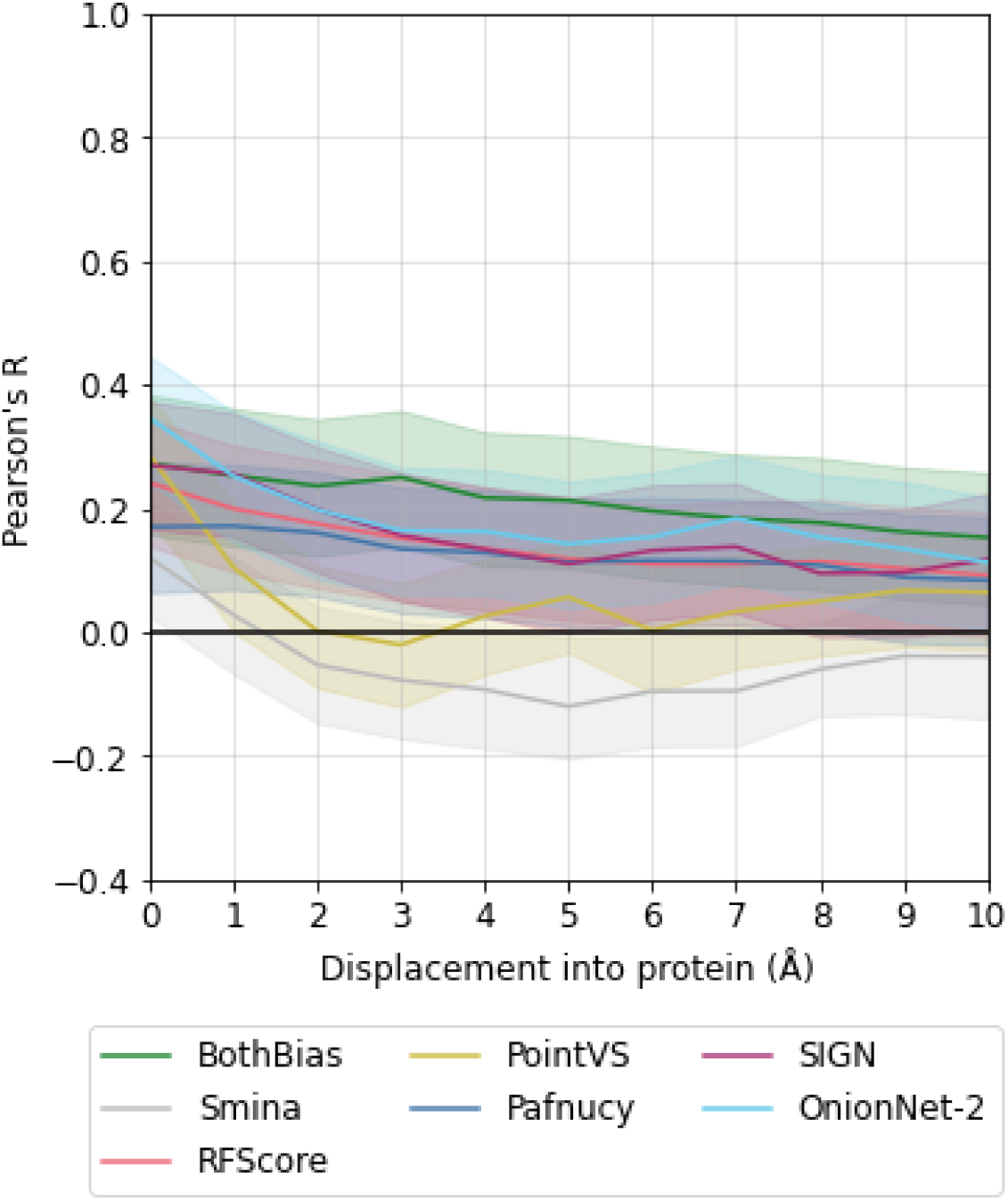
Pearson’s R between predicted and true pK values for protein-ligand complexes for our baseline models (LigandBias, ProteinBias and BothBias), a non-ML-based scoring function (Smina) and five commonly used MLBSFs (RFScore, PointVS, Pafnucy, SIGN and OnionNet-2), on progressively displaced ligands into the protein originally from 0 Ligand Bias crystal structures. Errors are the 95% confidence intervals from the bootstrapped Pearson’s R (N=10000).

